# Zinc deficiency decreases bone mineral density of rat by calmodulin-induced change in calcium metabolism

**DOI:** 10.1101/2020.06.09.143396

**Authors:** Qingli Yu, Jiali Zhao, Yanfeng Chen, Zixiang Li, Yongzhi Sun, Lina Fan, Maoqing Wang, Chenghai Peng

## Abstract

**Background:** Zinc deficiency decreases bone mineral density (BMD), but it is not known whether decreased BMD is a result of altered calcium absorption, excretion, and/or tissue distribution. To identified the associations between zinc deficiency, calcium metabolism and decreased BMD.

**Methods:** we performed two zinc deficiency experiments. In the first experiment, male rats (5-week-old) were fed a low zinc diet for four weeks. We measured serum zinc, alkaline phosphatase, Ca2+, osteocalcin, calcitonin, parathyroid hormone (PTH), calcium concentrations in feces and urine, BMD, and femur bone length and weight. In the second experiment, male rats (3-week-old) were fed a low zinc diet for five weeks. In addition to the aforesaid indicators, we measured the concentrations of zinc, total calcium, and calmodulin in multiple tissues.

**Results:** In both experiments, serum zinc, alkaline phosphatase, fecal and urine calcium, BMD, and bone weight of the low zinc diet group (LZG) were reduced compared with the normal zinc diet group (NZG)and pair-fed group (PZG); PTH increased significantly. Serum Ca2+, osteocalcin, and calcitonin concentrations were unchanged and not associated with decreased BMD. In the second experiment, zinc concentrations were reduced in serum, skeletal muscle, feces, and urine of LZG animals compared with NZG and PZG. Calmodulin in serum and skeletal muscle of the LZG group was decreased. Zinc deficiency increased total calcium concentrations in serum and skeletal muscle by promoting a decrease in calmodulin. To maintain blood Ca2+ balance, elevated PTH increased calcium reabsorption, reduced calcium excretion, stimulated bone resorption, mobilized bone calcium, and decreased BMD. (4) Conclusions: Decreased calmodulin and increased PTH induced by zinc deficiency altered calcium tissue distribution and decreased BMD of rat.

## 1. Introduction

Zinc is an essential element for growth of human and animals. Zinc is also an important mineral for the growth, development, and maintenance of bone and bone mineral density (BMD) [1–3]. Bone growth retardation and decreased BMD have been associated with zinc deficiency [4–7] in growing animals [8,9] and in infants[10], children and adolescents[5,11,12]. Zinc functions in bone metabolism by stimulating osteoblastic bone formation and inhibiting osteoclastic bone resorption [13,14]. Zinc supplementation prevents the bone loss and decreased BMD induced by various bone disorders [1,15–17]. However, it is not known mechanistically how zinc deficiency causes decreased BMD.

Calcium is directly related to bone development and metabolism. Adequate calcium intake is essential for the maintenance of bone health and normal BMD. Chronic calcium deficiency is a key cause of osteopenia and osteoporosis (decreased BMD). Calcium and zinc have antagonistic actions in a variety of systems [18]. Calcium is required for many types of cellular activation, whereas zinc tends to be a cellular inhibitor. Calmodulin is a ubiquitous regulator involved in many calcium-mediated processes. Calmodulin is activated by calcium and inhibited by zinc deficiency [19,20]. Calmodulin inhibition by zinc deficiency provides a rational molecular mechanism for zinc’s antagonism of calcium effects [21,22]. A defect in calcium channels was the first limiting biochemical defect found for zinc deficiency [23,24]. Zinc deficiency decreases calcium excretion [25]and increases calcium retention[26], which, in turn, increases calcium concentrations in blood cells, platelets, and tissues. Therefore, zinc is required for normal calcium metabolism[27]. In view of this fact, we speculated that decreased BMD is closely related to changes in calcium and calmodulin induced by zinc deficiency. This proposed relationship has not been investigated.

To identified the associations between zinc deficiency, calcium metabolism and decreased BMD, we performed two low zinc diet experiments with rats. First, we measured the concentrations of calcium in serum, feces, and urine, and quantitated key enzymes and hormones involved in calcium metabolism, such as parathyroid hormone (PTH), alkaline phosphatase (ALP), osteocalcin, calcitonin, and calmodulin. Then, we assessed overall calcium metabolism in zinc-deficient rats, including absorption, excretion, mobilization, and distribution, to explain the function of zinc in decreased BMD. We found that zinc deficiency decreased calmodulin, it increased PTH, which altered calcium metabolism, especially calcium tissue distribution, and zinc deficiency decreased BMD.

## 2. Materials and Methods

### Animal experiments

For the first experiment, 45 male Wistar rats (5-week-old, 110-130 g) were purchased from Beijing Vital River Laboratory Animal Technology Co., Ltd and housed individually with a 12-h light/dark cycle. The room temperature was controlled at 24 ± 1 °C and the relative humidity was 50% ± 5%. After one week of adaptive feeding, the rats were randomly divided into three groups: normal zinc diet group, NZG, low zinc diet group, LZG, and pair-fed zinc group, PZG. The standard diet was based on the AIN-93G diet (American Institute of Nutrition) with dried egg white as the protein source from Beijing KeAoLiXie Animal Food co., LTD, China[26]. The zinc content in the normal and low zinc diet was 30 and 10 mg/kg, respectively. PZG animals were fed with normal zinc diet (30 mg/kg zinc) and the food intake was the mean intake of the LZG. Body weight and food intake were measured at regular intervals. After the 4-week feeding period, all rats were fasted for 12 hours. Then, the animals were anesthetized by intraperitoneal injection of 10% chloral hydrate solution (0.3 ml/kg body weight). Blood was collected from the abdominal aorta and allowed to stand at room temperature for 2 hours. After centrifugation at (835 x g) for 10 minutes, the serum was separated and frozen at −80 °C. Urine and fecal samples were collected in metabolic cages with a sharp bottom funnel (see reference[26] for collection details). The left femur was dissected from muscle tissue, and femoral lengths and weights were measured.

In a second experiment, 3-week-old male rats (n=45) were fed for five weeks, and all other conditions were the same as in the first experiment. In addition to the foregoing samples, heart, skeletal muscle, kidney, and testicles were collected, washed with isotonic saline, weighed, and stored at −80 °C. The Ethics Committee of Harbin Medical University approved these animal experiments.

### Determination of serum biochemical indicators, calcium, calmodulin, and BMD

Ionized zinc and calcium in serum were measured by a colorimetric method (Sigma-Aldrich Trading Co., Ltd., Shanghai). BMD was measured with a dual X-ray energy bone densitometer (Norland XR-36 DEXA System, Cooper Surgical, Trumball, CT, USA) in the Second Affiliated Hospital of Harbin Medical University China.

ELISA assays (Summers Biomedical Products Co., Ltd., Beijing) were used to measure PTH, ALP, osteocalcin, calcitonin, and calmodulin in serum and tissues. Inductively coupled plasma mass spectrometry (ICP-MS) was used to measure total calcium and zinc concentrations in serum, feces, urine, liver, heart, skeletal muscle, kidney, and testis, as described[26].

### Statistical Analysis

All data were expressed as mean ± SD. Differences between groups were analyzed by an independent t test and ANOVA test using SPSS software (version 17.0, Chicago, IL, USA). A two-tailed value of P<0.05 was considered statistically significant.

## 3. Results

### Body weight and food intake

As shown in Figure 1A and 1B, body weights of LZG animals from the third week in the first experiment and from the second week in the second experiment were lower than weights of NZG and PZG animals (P <0.05). Food intake of LZG was significantly lower than NZG and PZG (P <0.05). In both experiments, there were significant decreases in serum zinc concentration in LZG compared with NZG and PZG (P< 0.05). We did not detect any significant differences in serum zinc between NZG and PZG in either experiment (P> 0.05).

**Figure-1.**
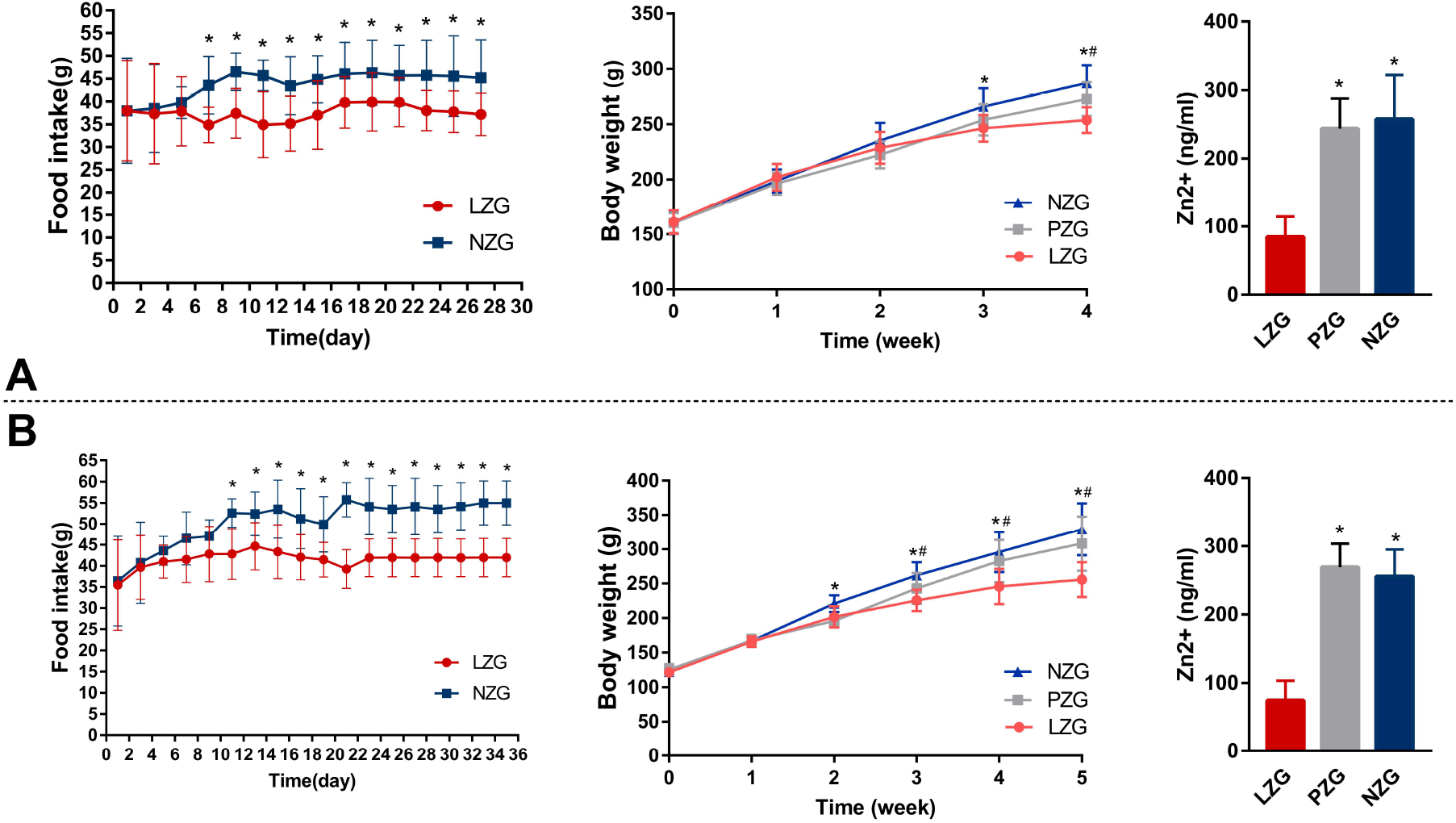
Food intake, body weight, and serum zinc concentration in two experiments. A: the first experiment, B: the second experiment. *P<0.05, LZG vs NZG and PZG, #, P<0.05, NZG vs PZG

### Femoral BMD, length, weight and serum Ca2+ concentration

In both experiments, femoral BMD and weights of LZG were lower than those of NZG and PZG (P<0.05; Figure 2). The femoral lengths in the first experiment were not statistically different between the three groups (P>0.05), whereas the femoral lengths of LZG were significantly shorter than those of NZG and PZG in the second experiment (P<0.05). We did not find any significant differences in BMD, femoral length, or weight between NZG and PZG in both experiments (P> 0.05). Lastly, there were no statistical differences in serum Ca2+ concentrations between the three groups in both experiments (P>0.05; Figure 2).

**Figure-2.**
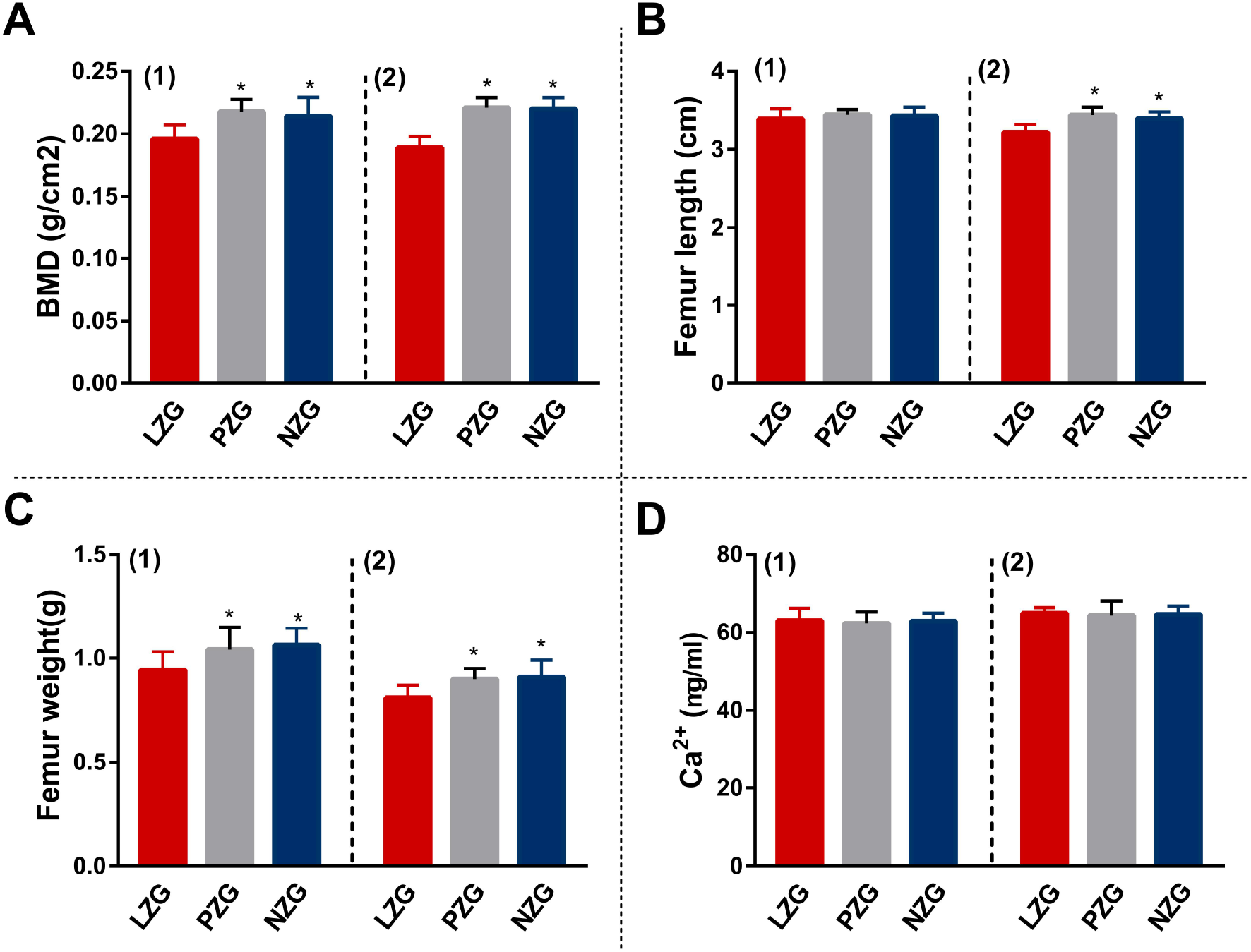
Femoral length, weight, BMD and serum Ca2+ concentration in two experiments. (1): the first experiment, (2): the second experiment. A: BMD, B: femur length, C: femur weight, D: Ca2+. *P<0.05, LZG vs NZG and PZG

### Key enzymes and hormones involved in calcium metabolism

The concentrations of decreased ALP (Figure 3A) and increased PTH (Figure 3B) were observed (P < 0.05) in LZG compared with normal zinc diet groups (P <0.05) in both experiments. We did not detect any significant differences in osteocalcin (Figure 3C) and calcitonin (Figure 3D) between the three groups(P>0.05).

**Figure-3.**
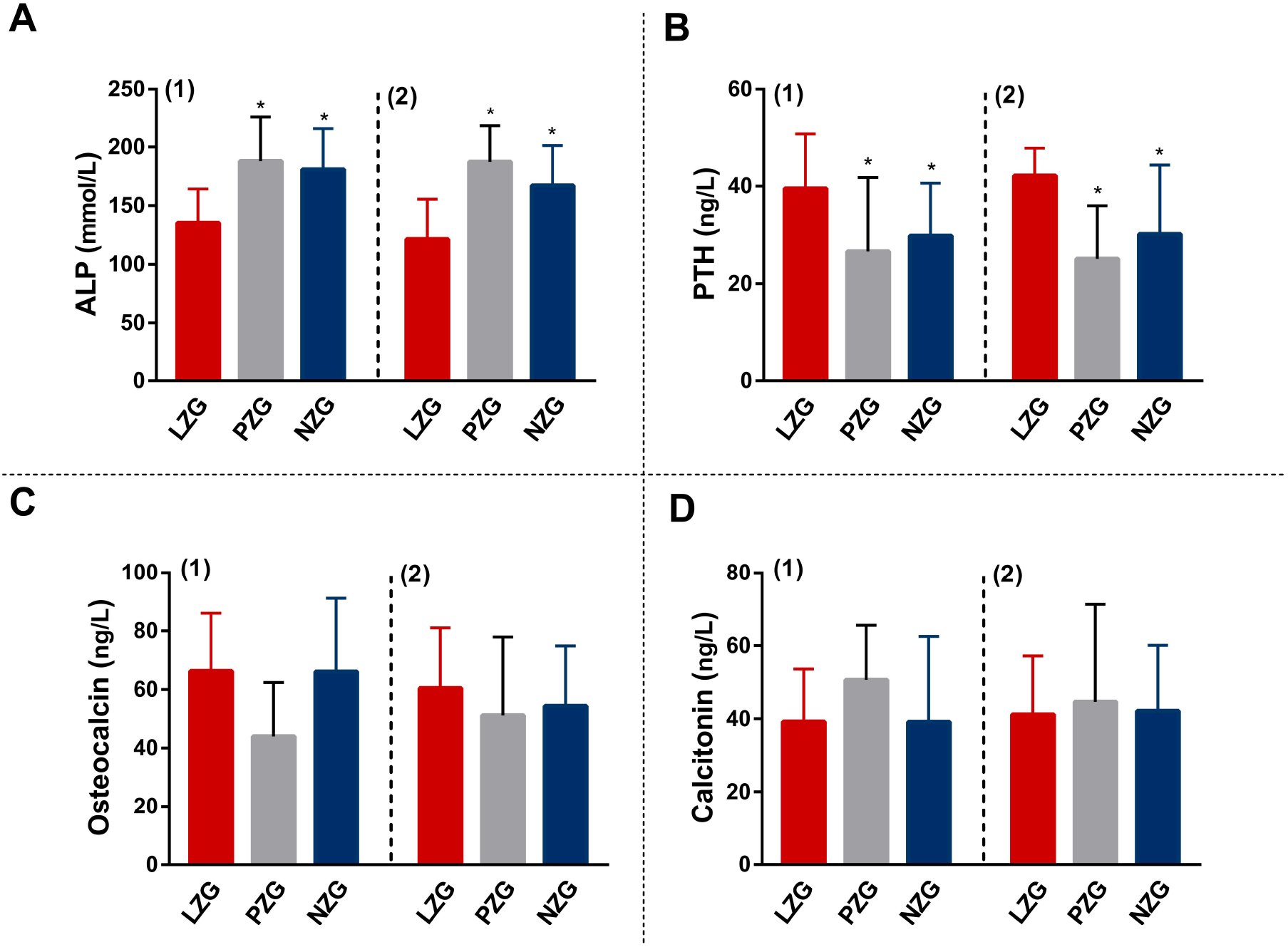
Concentrations of ALP, PTH, osteocalcin and calcitonin in serum in two experiments. (1): the first experiment, (2): the second experiment, A: ALP, B: PTH, C: osteocalcin, D: calcitonin, *P<0.05, LZG vs NZG and PZG

### Total calcium concentrations in serum, feces, urine and multi tissues

In the first experiment, we found decreases in total calcium in feces, urine, and liver, and increases in total calcium in serum in LZG compared with NZG and PZG (P<0.05) in our published article[26]. We did not collect other tissues in this first experiment, thus, the concentrations of calcium in other tissues were unknown.

In the second experiment, we measured decreases in calcium in feces, urine, and liver, and increased total calcium in serum and skeletal muscle in LZG compared with NZG and PZG (P<0.05, Figure 4). There were no statistical differences in total calcium in heart, kidney, and testis between the three groups (Sup Figure 1). There were no differences in total calcium of serum, urine, feces, and all tissues between NZG and PZG (P> 0.05; Figure 4A and Sup Figure 1). Calmodulin concentrations in serum and skeletal muscle of LZG were decreased compared with NZG and PZG (P<0.05, Figure 4B), whereas there were no differences in other tissues (Sup Figure 1B). Calmodulin in all tissues was not significantly different between NZG and PZG (P> 0.05).

**Figure-4.**
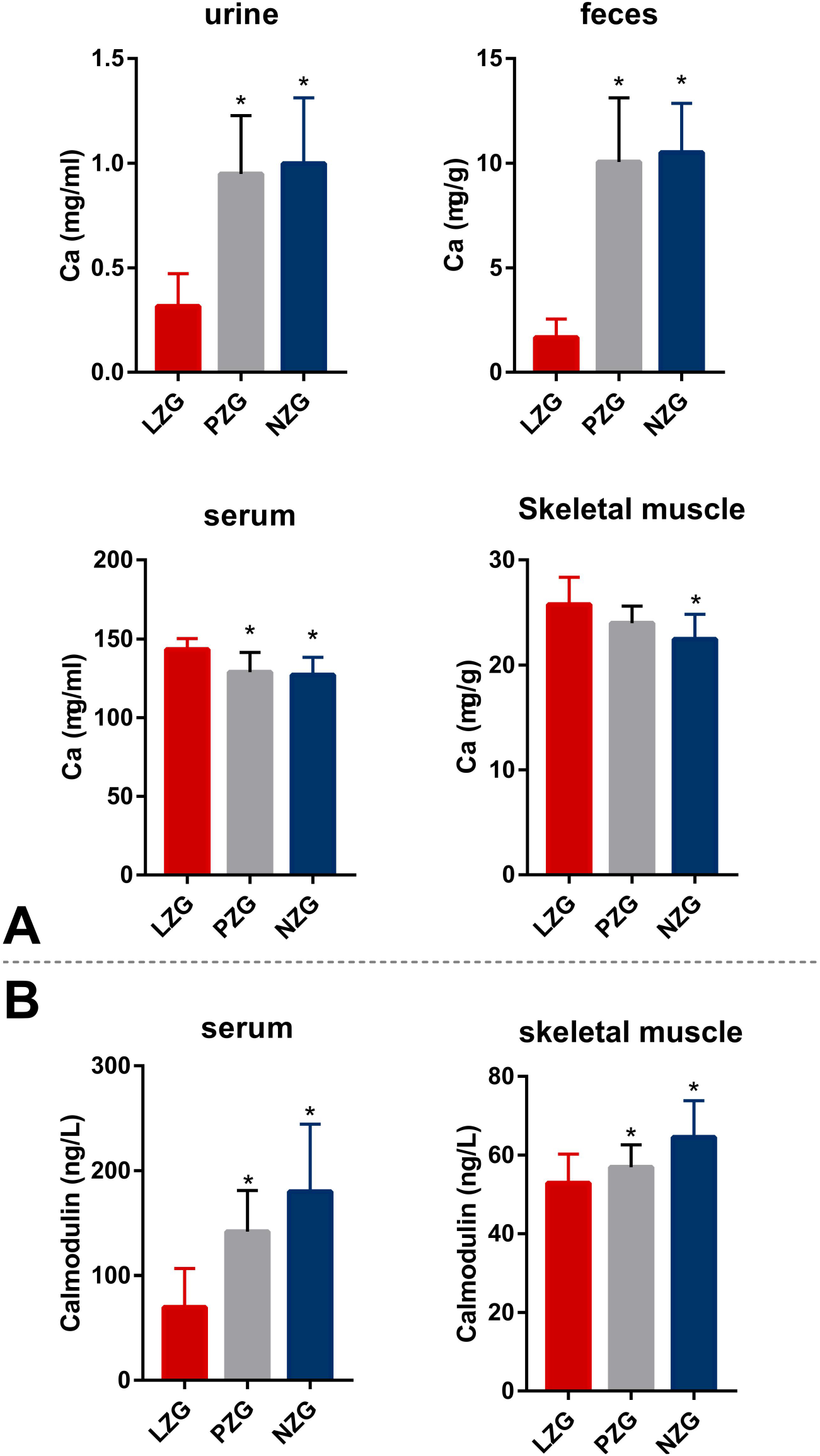
Concentrations of total calcium and calmodulin in different tissues in the second experiment. A: Total Ca concentrations in different tissues, B: calmodulin concentrations. *P<0.05, LZG vs NZG and PZG

## 4. Discussion

Calcium is directly related to skeletal development, and zinc deficiency causes decreased BMD. However, it was not known whether zinc deficiency decreases BMD by altering calcium absorption, excretion, and/or tissue distribution. By measuring the concentrations of calcium in serum, feces, urine, and various tissues, and the concentrations of calmodulin in various tissues, we identified the associations between low Zn intake and Ca metabolism and decreased BMD.

In both of our animal experiments, zinc deficiency all decreased BMD and femur bone weight (Figure 2). In the second experiment, due to the relatively young age of the rats and a longer feeding period, the low zinc diet affected bone development and decreased femur length. In our previous study, we found that a low calcium diet (12 weeks) decreased rat BMD[28]. However, in the present study, we found that a low zinc diet (four weeks) decreased BMD and femur length of LZG (Figure 2). Thus, zinc deficiency had a greater negative effect on BMD compared with calcium deficiency. Therefore, it was necessary to reveal the relationship between zinc deficiency and decreased BMD.

### Relationship between zinc deficiency, calcium metabolism and decreased BMD

In the two experiments, we measured significant increases in PTH and decreases in ALP in LZG; the serum concentrations of calcitonin and osteocalcin were unchanged between the three groups (Figure 3). Thus, significantly increased PTH and decreased ALP were associated with decreased BMD, and calcitonin and osteocalcin were not associated with decreased BMD.

In the first experiment, we did not find changes in serum Ca2+ (Figure 2), but calcium level and calcium excretion in feces and urine of LZG (Figure 4) were reduced compared with NZG and PZG. These changes may have been related to elevated PTH. Elevated PTH increases reabsorption of calcium, decreases calcium level, and decreases calcium excretion in urine and feces. Our results suggested that zinc deficiency increased calcium reabsorption and calcium of LZG was not lost[26]. Serum Ca2+ was also not different between the three groups in the second experiments (Figure 2). There was no difference in dietary calcium intake between LZG and PZG (Figure 1). Thus, why did the BMD of LZG decrease, and where was the calcium that was lost from the femurs of LZG? Other investigators have found that zinc deficiency increased calcium retention in other tissues [19]. Therefore, we speculated that zinc deficiency increased the distribution of calcium in serum or other tissues by the combined effects of calcium-activated calmodulin and decreased BMD.

We tested the foregoing hypothesis in the second experiment. In addition to serum, urine, and feces, we collected various tissues and measured their concentrations of total calcium, zinc, and calmodulin to reveal the effect of zinc deficiency on calcium metabolism. We also observed decreased calcium in feces and urine of LZG. This result confirmed that calcium was not lost in zinc-deficient rats. As we expected, we found increased total calcium in serum and skeletal muscle in LZG compared with NZG and PZG (Figure 4 A), and there were no changes in other tissues, including heart, kidney, and testis (Sup Figure 1). These results further confirmed our speculation that zinc deficiency redistributed calcium and increased the concentrations of total calcium in serum and skeletal muscle. The calcium redistribution of LZG correlated with decreased BMD.

Zinc and calcium exhibit antagonistic actions in metabolism. Zinc deficiency inhibits calmodulin activity, whereas calcium increases calmodulin activity. In the second experiment, decreased zinc (Sup Table 1) and decreased calmodulin (Figure 4B) were both observed in LZG compared with NZG. Without sufficient zinc, more calcium in blood and skeletal muscle was needed to increase or maintain the activity of calmodulin. To increase calcium in serum and skeletal muscle, elevated PTH promoted calcium reabsorption and inhibited calcium excretion in urine and feces (Figure 5). Therefore, compared with NZG, we observed decreased calcium concentrations and excretion in urine and feces of LZG in both experiments (Figure 4).

**Figure 5.**
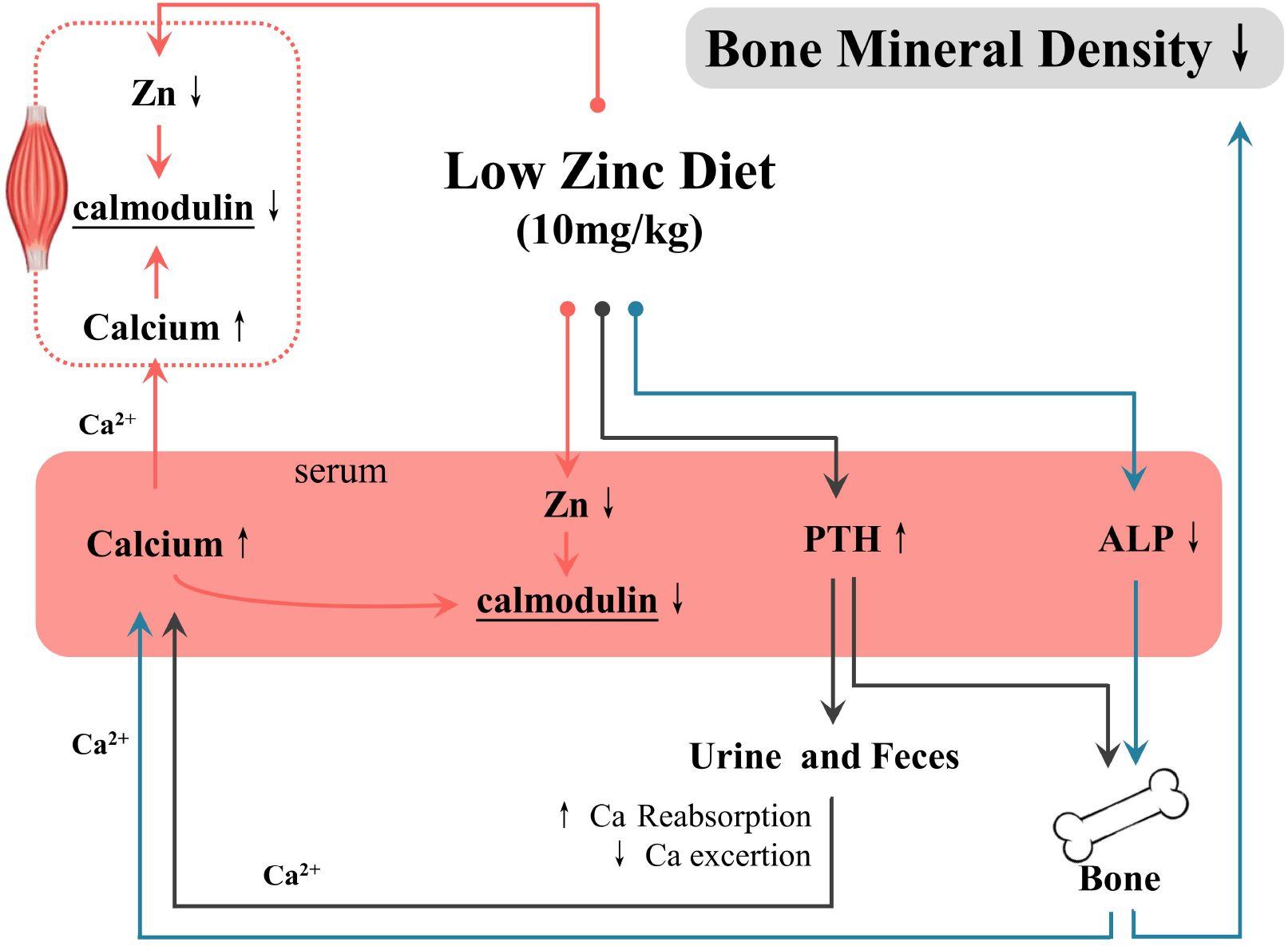
Relationship between zinc deficiency, altered calcium metabolism and decreased BMD. ↑: LZG vs NZG and PZG; ↓: LZG vs NZG and PZG

Calcium concentrations in the blood and fluid that surround cells are tightly controlled to preserve normal physiological function. The skeleton is a reserve of calcium which is drawn upon to maintain adequate serum calcium when calcium intake is limited. Elevated PTH stimulates bone resorption and release of bone calcium for restoring serum calcium concentration, a process that ultimately causes a decrease in BMD[29] (Figure 5). If LZG had continued to be fed a low zinc diet, the calcium deficiency of rat would have been more severe. But, we can not explain the reason of decreased Ca in liver in our study.

Several studies found that serum zinc was associated with BMD among children 0-5 years-old and adolescent girls[5,11,12]. On the other hands, zinc supplementation can improve bone density of young adults[30]. However, the above studies did not clearly reveal the relationship between zinc deficiency and decreased BMD in children and adolescents. Our study provided a foundation for tackling the mechanism of the low zinc mediated impairment of normal Ca metabolism. Therefore, we recommend that children and adolescents should monitor the nutritional status of zinc because zinc helps to resist the shift in distribution of calcium and maintains their bone health.

## 5. Conclusions

Zinc deficiency reduced calmodulin in serum and skeletal muscle and increased total calcium concentrations in these sources. To maintain blood calcium balance, elevated PTH increased calcium absorption, decreased excretion, and mobilized bone calcium, eventually leading to decreased BMD of rat. Children and adolescents must ensure proper zinc intake for maintaining their bone health.

## Supplementary Materials

**Supplemental Table-1.** The concentrations of Zinc in 8 tissues of rats.

|  | LZG | PZG | NZG |
| --- | --- | --- | --- |
| Feces (µg/g) | 23.23±6.03 | 247.68±19.79* | 253.31±36.17* |
| Urine (µg/ml) | 4.56±1.21 | 9.34±2.66* | 8.16±2.72* |
| Serum (µg/ml) | 1.89±0.09 | 2.26±0.33* | 2.19±0.44* |
| Skeletal muscle (µg/g) | 14.82±2.08 | 15.61±1.62 | 18.72±3.49* |
| Liver (µg/g) | 47.26±2.89 | 57.70±4.65* | 60.22±5.46* |
| Kidney (µg/g) | 23.10±1.55 | 25.55±3.33 | 27.15±2.72 |
| Testis (µg/g) | 19.77±2.46 | 10.75±1.89* | 13.84±2.44* |
| Heart (μg/g) | 19.68±1.50 | 18.86±1.79 | 20.43±1.25 |

\*: P<0.05 LZG vs NZG and PZG.

**Supplemental figure-1.**
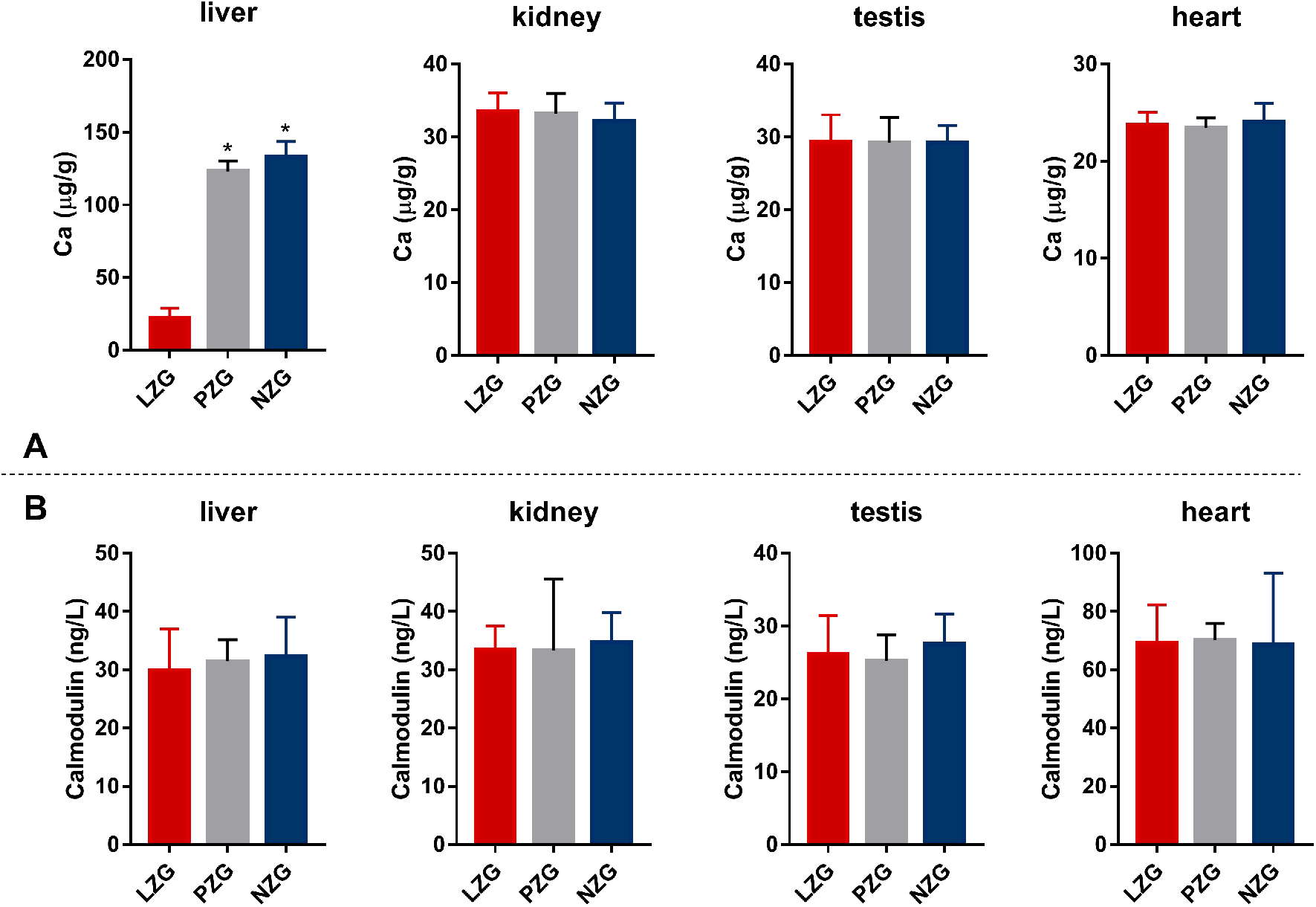
The concentrations of total calcium and calmodulin of different tissues in the second experiment. A: Total calcium, B: calmodulin.

## Author Contributions

Maoqing Wang and Chenghai Peng contributed to the study’s conception and design. Qingli Yu and Jiali Zhao drafted the manuscript. Qingli Yu performed data analysis and interpretation. Zixiang Li, Yongzhi Sun and Lina Fan performed animal experiments and data collection. Yanfeng Chen measured element concentrations by ICP-MS. The final draft was read and approved by all the authors.

## Acknowledgments

The authors thank AiMi Academic Services (www.aimieditor.com) for English language editing and review services.

